# A Distinct Contractile Injection System Found in a Majority of Adult Human Microbiomes

**DOI:** 10.1101/865204

**Authors:** Maria I. Rojas, Giselle S. Cavalcanti, Katelyn McNair, Sean Benler, Amanda T. Alker, Ana G. Cobián-Güemes, Melissa Giluso, Kyle Levi, Forest Rohwer, Sinem Beyhan, Robert A. Edwards, Nicholas J. Shikuma

## Abstract

An imbalance of normal bacterial groups such as Bacteroidales within the human gut is correlated with diseases like obesity. A current grand challenge in the microbiome field is to identify factors produced by normal microbiome bacteria that cause these observed health and disease correlations. While identifying factors like a bacterial injection system could provide a missing explanation for why Bacteroidales correlates with host health, no such factor has been identified to date. The lack of knowledge about these factors is a significant barrier to improving therapies like fecal transplants that promote a healthy microbiome. Here we show that a previously ill-defined Contractile Injection System is carried in the gut microbiome of 99% of individuals from the United States and Europe. This type of Contractile Injection System, we name here Bacteroidales Injection System (BIS), is related to the contractile tails of bacteriophage (viruses of bacteria) and have been described to mediate interactions between bacteria and diverse eukaryotes like amoeba, insects and tubeworms. Our findings that BIS are ubiquitous within adult human microbiomes suggest that they shape host health by mediating interactions between Bacteroidales bacteria and the human host or its microbiome.

## INTRODUCTION

The human gut harbors a dynamic and densely populated microbial community (Gill et al., 2006). The gut microbiome of healthy individuals is characterized by a microbial composition where members of the Bacteroidetes phylum (*Bacteroides* and *Parabacteroides*) constitute 20-80% of the total (Consortium et al., 2012). Several studies have demonstrated that dysbiosis in the human gut is correlated with microbiome immaturity, and diseases like obesity and inflammatory bowel disease (Carroll et al., 2011; Hooper and Gordon, 2001; Kau et al., 2011; Ley et al., 2006; Subramanian et al., 2014). However, we currently do not know the identity of bacterial factors that explain these correlations between Bacteroidales abundances and healthy human phenotypes. The lack of knowledge about these factors is a significant barrier to improving therapies like fecal transplants that manipulate microbial communities to treat microbiome-related illness.

Many bacteria such as Bacteroidales produce syringe-like secretion systems called Contractile Injection Systems (CIS) that are related to the contractile tails of bacteriophage (bacterial viruses) (Cianfanelli et al., 2016; Salmond and Fineran, 2015). Most CIS characterized to date, termed Type 6 Secretion Systems (T6SS), are produced and act from within an intact bacterial cell. In contrast, extracellular CIS (eCIS) are released by bacterial cell lysis, paralleling the mechanism used by tailed phages to escape their host cell (**Figure 1A**). To date, Bacteroidales from the human gut have only been shown to produce one type of CIS, a Subtype-3 T6SS that mediates bacteria-bacteria interactions and can govern the microbial composition of the gut microbiome (Chatzidaki-Livanis et al., 2016; Coyne et al., 2016; Russell et al., 2014; Verster et al., 2017). However, CIS from other bacterial groups such as Gammaproteobacteria are known to help bacteria interact with diverse eukaryotic organisms such as amoeba, insects, tubeworms and humans.

**Figure 1.**
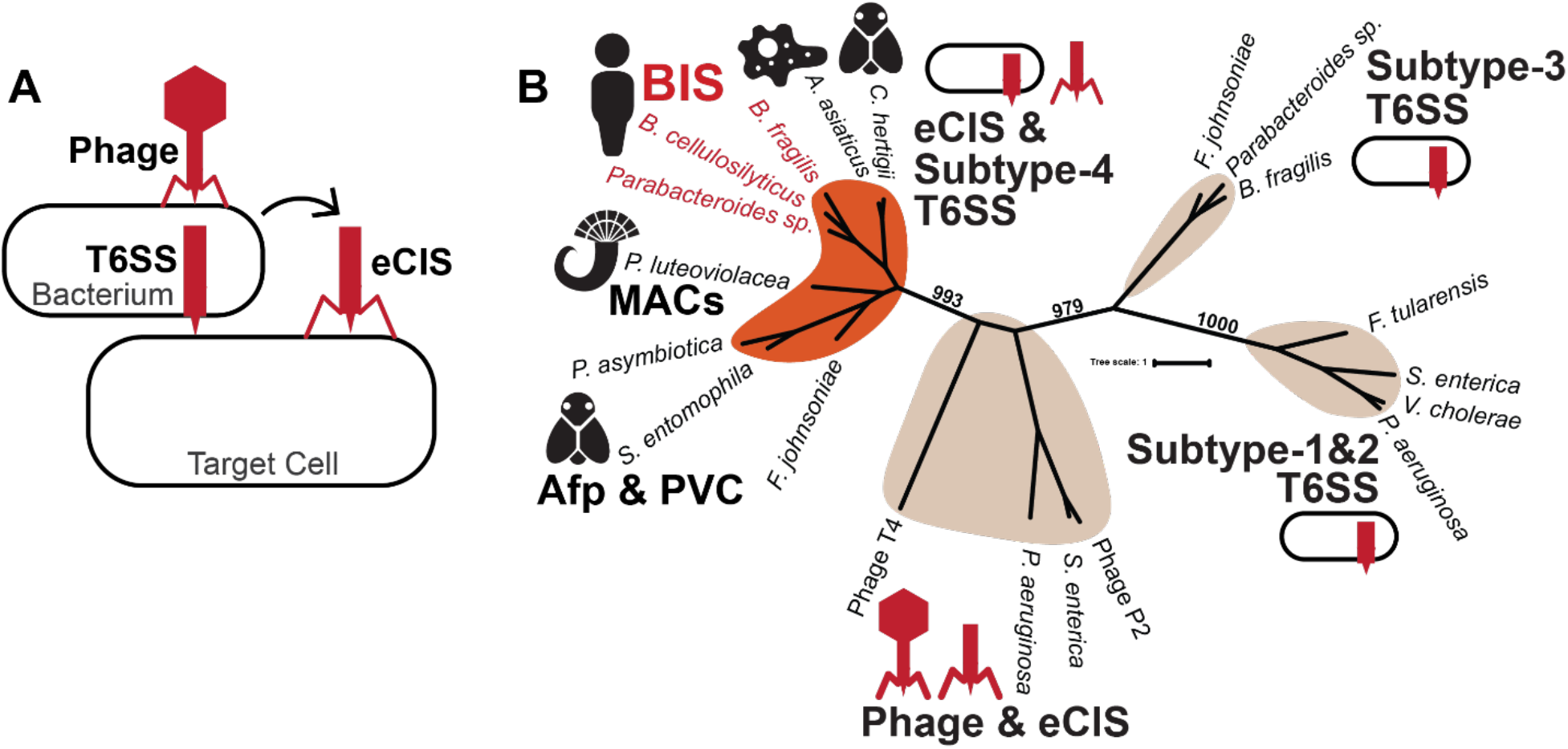
Bacteroidales possess a distinct Contractile Injection System. **(A)** Contractile Injection Systems are related to the contractile tails of bacteriophage. There are two main types of CIS; Type 6 Secretion Systems (T6SS) act from within a bacterial cell, while extracellular CIS (eCIS) are released by bacterial cell lysis and bind to target cells. **(B)** Unrooted phylogeny of CIS sheath protein sequences. BIS group with known Subtype-4 T6SS and eCIS (orange) and are distinct from known Subtype-1, Subtype-2 and Subtype-3 T6SS. Bacteria with BIS identified in this study are highlighted in red.

Distinct from Bacteroidales Subtype-3 T6SS is a different class of CIS that may have evolved independently (Böck et al., 2017). Intriguingly, all previously described examples of these distinct CIS mediate bacteria-eukaryote interactions. Three are classified as eCIS and include: (1) MACs (Metamorphosis Associated Contractile Structures) that stimulate the metamorphosis of tubeworms (Shikuma et al., 2016, 2014), (2) PVCs (*Photorhabdus* Virulence Cassettes) that mediate virulence in grass grubs (Yang et al., 2006), and (3) Afp (Anti-Feeding Prophage) that cause cessation of feeding and death of grass grub larvae (Heymann et al., 2013; Hurst et al., 2018, 2007, 2004). A fourth CIS from *Amoebophilus asiaticus* (Böck et al., 2017) promotes intracellular survival in amoeba and defines the Subtype-4 T6SS group. Although this class of CIS forms a superfamily of structures found in diverse microbes (Chen et al., 2019), until the present work, examples of this distinct class of CIS have only been identified in a few isolated Bacteroidetes genomes (Böck et al., 2017; Penz et al., 2012; Sarris et al., 2014).

In this study, we show that diverse Bacteroidales species from the human gut encode a distinct CIS within their genome that is related to eCIS and T6SS that mediate tubeworm, insect and amoeba interactions (MACs, PVCs, Afp, Subtype-4 T6SS). Strikingly, we show that this distinct CIS from Bacteroidales are present within the microbiomes of over 99% of human adult individuals from Western countries (Europe and the United States) and are expressed *in vivo*. Our results suggest that a previously enigmatic class of contractile injection system is present and expressed within the microbiomes of nearly all adults from Western countries.

## RESULTS

### Bacteroidales bacteria from the human gut possess genes encoding a distinct Contractile Injection System

Using PSI-BLAST to compare previously identified eCIS and Subtype-4 T6SS proteins to the non-redundant (nr) protein sequence database, we identified CIS structural proteins (baseplate, sheath and tube) that matched with proteins from various human Bacteroidales isolates, including a bacterial isolate from the human gut (*Bacteroides cellulosilyticus* WH2, **Table S1**) (McNulty et al., 2013). To determine the relatedness of these distinct CIS with all known CIS subtypes, we performed phylogenetic analyses of CIS proteins that are key structural components of CIS—the CIS sheath and tube. Multiple methods of phylogenic analyses (maximum likelihood, neighbor joining, maximum parsimony, UPGMA, and minimum evolution) showed that Bacteroidales sheath and tube proteins consistently formed a monophyletic group with other eCIS and Subtype-4 T6SS sheath and tube proteins (**Figures 1B, S1, Table S2**). Moreover, the BIS sheath and tube were clearly distinct when compared to previously characterized *Bacteroides* Subtype-3 T6SS (**Figures 1B, S1**) (Chatzidaki-Livanis et al., 2016; Russell et al., 2014). Based on these data and results below, we name this distinct CIS as Bacteroidales Injection Systems (BIS).

### Genes encoding BIS are found in a conserved cluster of genes and form three genetic arrangements

To identify Bacteroidales species that possess a bonafide BIS gene cluster, we performed a comprehensive search of 759 sequenced *Bacteroides* and *Parabacteroides* genomes from the Refseq database. Our sequence-profile search revealed 66 genomes from *Bacteroides* and *Parabacteroides* species that harbor complete BIS gene clusters **(Table S3)** in three conserved gene arrangements (**Figure 2A**). The first architecture is exemplified by *B. cellulosilyticus* WH2, which harbors two sheath proteins, two tube proteins, and a protein with unknown function intervening between putative genes encoding the baseplate (gp25, gp27 and gp6). The second architecture is exemplified by *B. fragilis* BE1. This architecture has a single sheath protein and lacks the hypothetical proteins observed in architecture one between gp25 and gp27, and between Tube2 and LysM. Finally, the third architecture defined by *P. distasonis* D25 is the most compact, lacks four hypothetical proteins found in architectures 1 and 2. Additionally, gp27 and gp6 proteins are shorter, and the genes FtsH/ATPase and DUF4157 are inverted. Importantly, all three genetic architectures have genes with significant sequence similarity to MAC, Afp and PVC genes shown previously to produce a functional CIS, including baseplate proteins (gp25, gp27, and gp6), sheath, tube, and FtsH/ATPase (**Figure 2B**).

**Figure 2.**
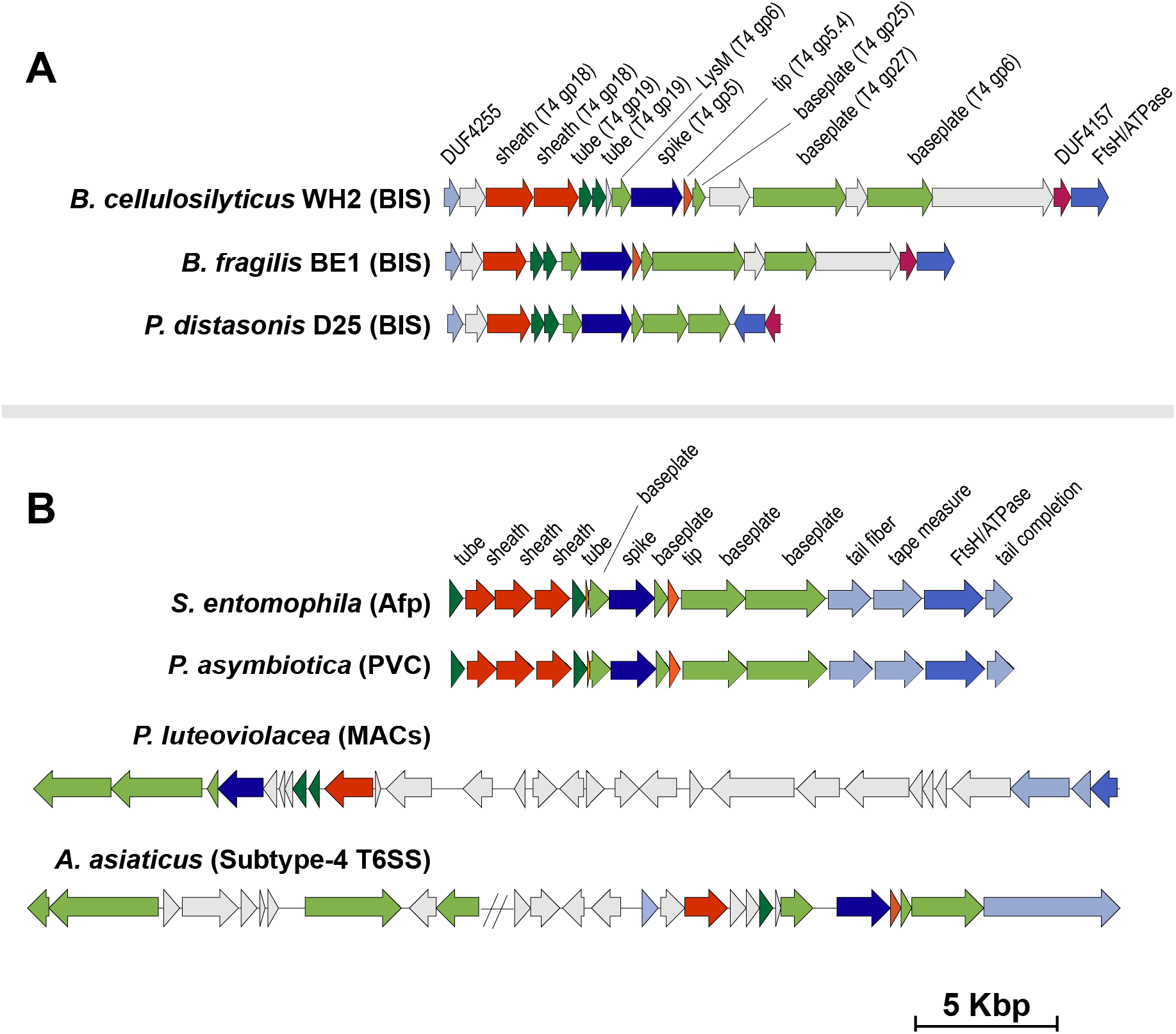
Bacteroidales produce three architectures of BIS. **(A)** Synteny plot of BIS gene clusters in *Bacteroides* and *Parabacteroides* species compared to **(B)** those of *P. luteoviolacea* MACs, *S. entomophilia* Afp, *Photorhabdus* PVCs, and *A. asiaticus* Subtype-4 T6SS. Representative CIS gene cluster architectures are shown, with genes color coded according to function. Genes with no significant sequence similarity at the amino acid level to any characterized proteins are colored light grey. Sequence coordinates of all gene clusters are provided in **Table S3**.

### BIS genes are present in human gut, mouth, and nose microbiomes

To determine the prevalence and distribution of BIS genes in human microbiomes, we searched shotgun DNA sequencing data from 11,219 microbiomes from the Human Microbiome Project database, taken from several locations on the human body (Consortium et al., 2012; Levi et al., 2018). We sampled these metagenomes for the presence of 18 predicted BIS proteins **(Table S4)**. Across all HMP metagenomes, 8,320 (74%) showed hits to at least one of the 18 BIS proteins. Hits were distributed across metagenomes from various mucosal tissues, and were more abundant in the gut and in the mouth, where Bacteroidales are found in high abundance (Consortium et al., 2012). The dataset included stool (1,851, 99.6% of total stool metagenomes), oral (4,739, 79.2% of total oral metagenomes), nasal (630, 41.8% of total nasal metagenomes), and vaginal (232, 27.7% of total vaginal metagenomes) samples (**Figure 3A**).

**Figure 3.**
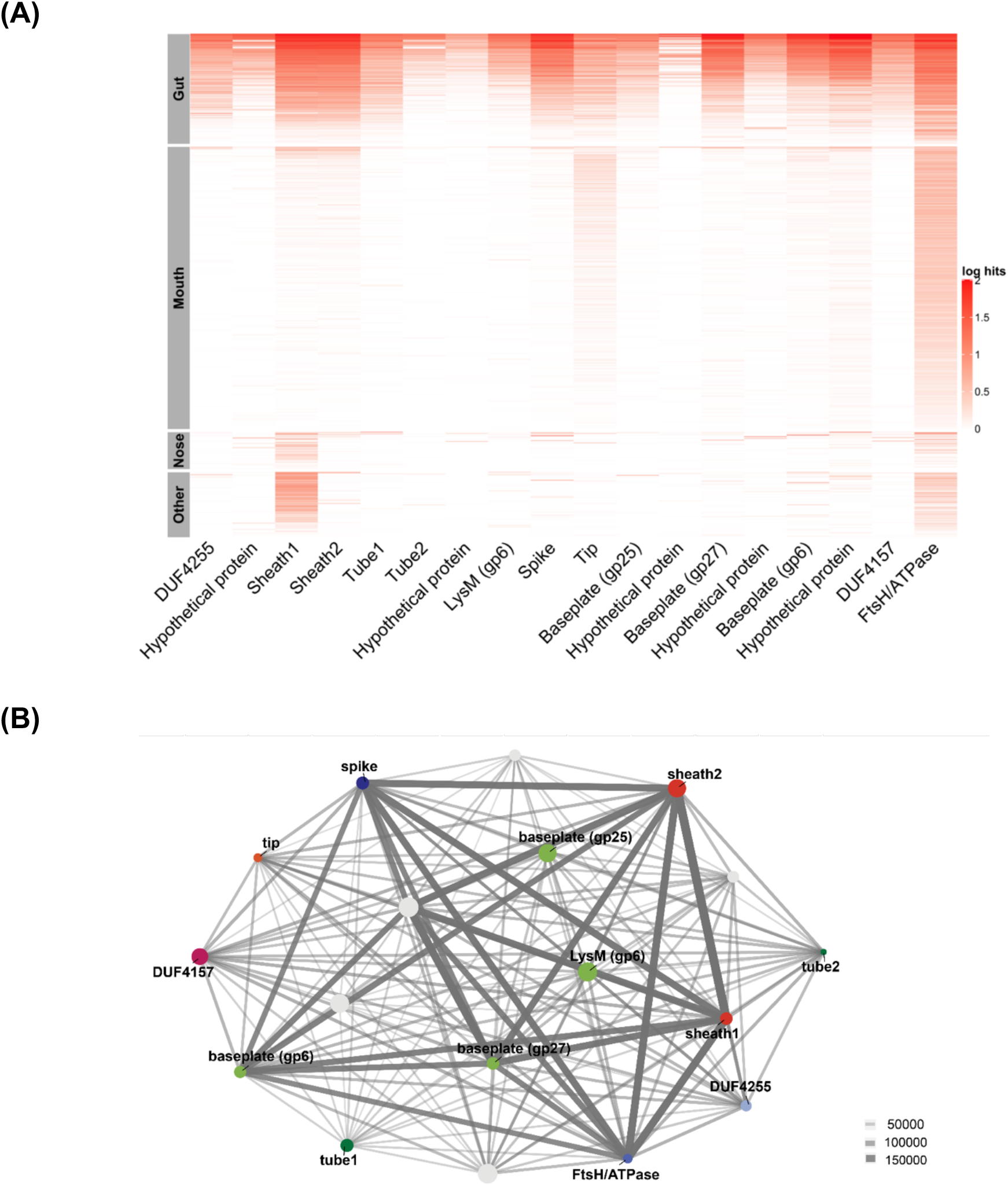
BIS genes are abundant in human gut and mouth microbiomes. **(A)** Coverage plot of BIS genes (Log10 of 1,000,000*hits/reads) in 8,320 microbiomes associated with mucosal tissue: gut, mouth, nose, and other (includes vaginal and skin tissues) from 300 healthy humans. **(B)** Ten BIS genes are found more often together in human metagenomes (co-occurrence network). Node size represents the number of hits for each protein across all runs. Line weight represents the number of times any two proteins occurred together within a dataset.

To determine how often any of the 18 genes co-occurred within the same metagenome sample, we constructed a co-occurrence network. Ten genes appeared together at high frequencies including Sheath1, Sheath2, FtsH/ATPase, baseplate (gp25, gp27, and gp6), LysM, Spike, and two hypothetical proteins (**Figure 3B**). The gene with the highest hit abundance encodes for an ATPase homologous to the *Escherichia coli* FtsH, known to be involved in cleavage of the lambda prophage repressor, followed by a hypothetical protein, and Sheath1. The other genes, including Tube1, Tube2, Tip, DUF4255 domain-containing protein, DUF4157 domain-containing protein, and three hypothetical proteins, were detected together less often within the microbiome samples.

### BIS genes are expressed *in vivo* in the gut of humanized mice, and *in vitro* when cultured with various polysaccharides

To determine whether BIS genes are transcribed under laboratory growth conditions or within the human gut, we searched for the 18 major BIS proteins in publicly available RNA sequencing data from *in vivo* metatranscriptomes of humanized mice (Ridaura et al., 2013) (gnotobiotic mice colonized with human microbiome bacteria) and *in vitro B. cellulosilyticus* WH2 pure cultures (McNulty et al., 2013). We inspected 59 metatranscriptomes from a previously published *in vivo* study (Ridaura et al., 2013) for the presence of the 18 major BIS proteins, where gnotobiotic mice were inoculated with human gut microbiome cultures. In 48 out of 59 metatranscriptomes (81.4%), we found expression of at least 15 BIS proteins **(Figure 4)**. Similarly, when *B. cellulosilyticus* WH2 was cultured in minimum media (MM) supplemented with 31 different simple and complex sugars (McNulty et al., 2013), all 18 genes were expressed at least once in at least two of the three replicate cultures (**Figure S2**). The highest expression was seen under growth in N-Acetyl-D-Galactosamine (GalNAc) and N-Acetyl-Glucosamine (GlcNAc), amino sugars that are common components of the bacterial peptidoglycan, in high abundance in the human colon, and implicated in many metabolic diseases (Baudoin and Issad, 2014; Myslicki et al., 2014). Our analyses of metatranscriptomes show that BIS genes are transcribed by Bacteroidales bacteria under laboratory growth conditions and within humanized mouse microbiomes *in vivo*.

**Figure 4.**
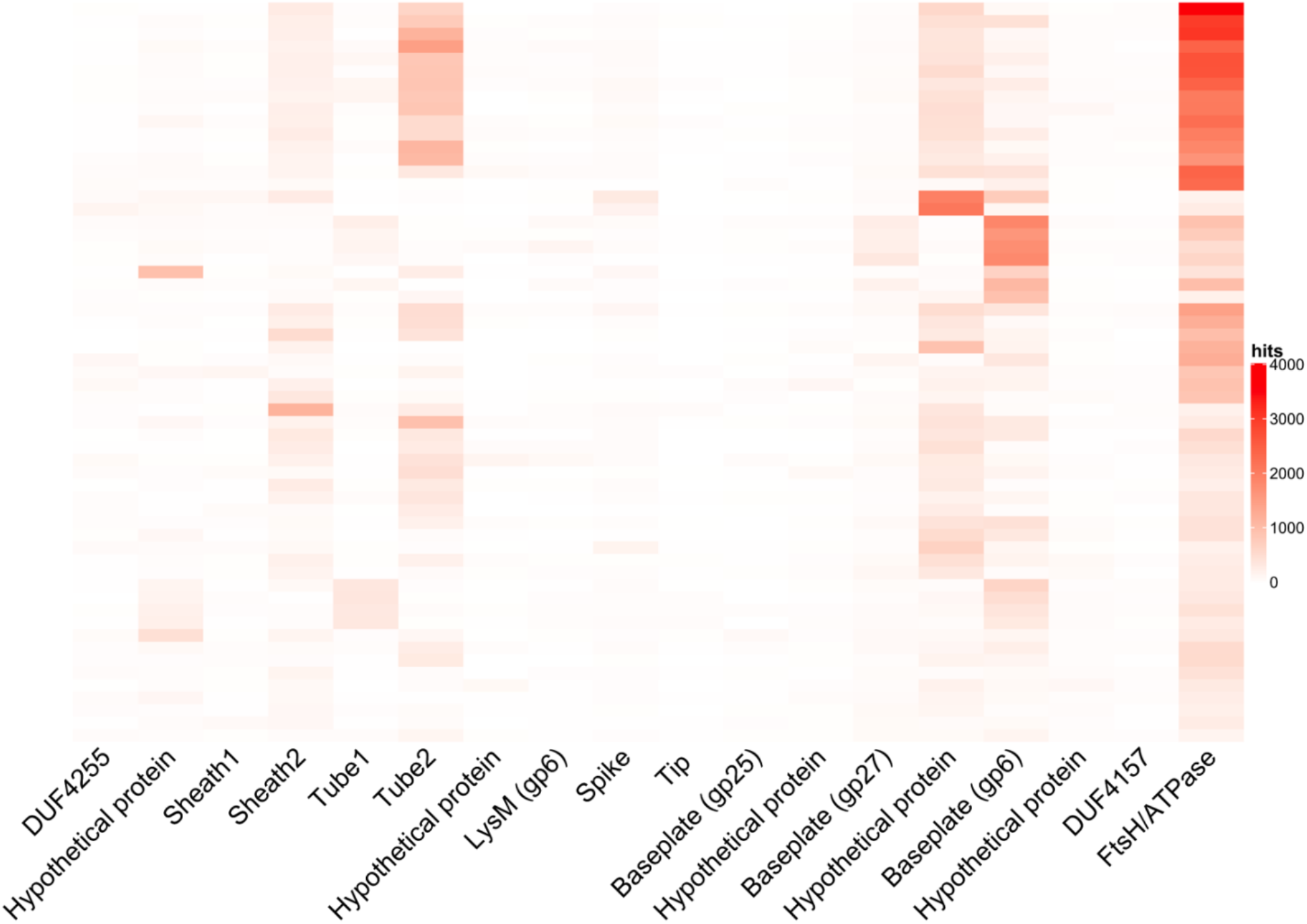
BIS genes are expressed *in vivo* in humanized mice. Coverage plot of BIS genes (normalized by number of reads and protein nt size) from stool metatranscriptomes of humanized mouse microbiomes.

### BIS genes are present in the microbiomes of nearly all adult individuals

To determine the prevalence of BIS within the microbiomes of human populations, we analyzed 2,125 fecal metagenomes from 339 individuals; 124 Individuals from Europe (Qin et al., 2010) and 215 individuals from North America (Consortium et al., 2012). We found that all individuals possessed at least 1 of the 18 BIS genes within their gut microbiome (**Figure 5**). Most individuals carried at least 9 BIS proteins (83.0% in HMP and 90.3% in Qin dataset). A lower number possessed all 18 BIS proteins (8.96% in HMP and 6.45% in Qin dataset).

**Figure 5.**
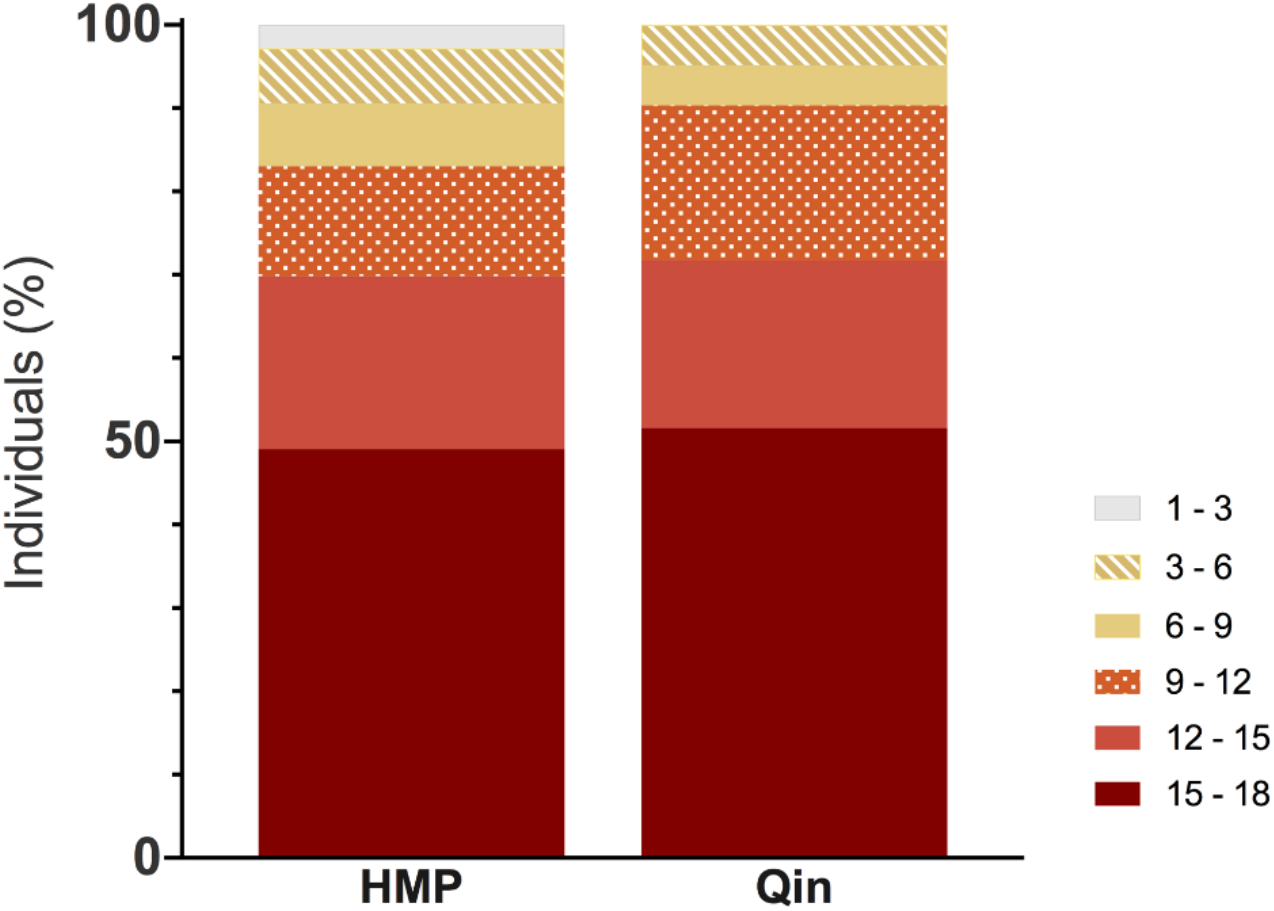
BIS genes are present in the microbiomes of a majority (99%) of adult individuals from the United States and Europe. Frequencies of 18 BIS proteins from fecal samples of 339 individuals. HMP n=215, from healthy North American individuals (Consortium et al., 2012); Qin n=124, from a study of European individuals (Qin et al., 2010). Protein hits are normalized by individual donor.

## DISCUSSION

Here, we show that a previously ill-described Contractile Injection System is present in the gut microbiomes of nearly all adult individuals from Western countries. We found that BIS genes are present in human microbiomes throughout mucosal tissues (oral, nasal, vaginal, ocular) and enriched in metagenomes from gut samples. CIS have gained recent recognition as secretion systems promoting disease in several prominent human pathogens like *Pseudomonas aeruginosa* and *Vibrio cholerae*. However, our discovery of a distinct class of CIS produced by Bacteroidales from human microbiomes stems from studies of symbiotic interactions between environmental bacteria and diverse eukaryotic hosts like tubeworms, insects and amoeba (Böck et al., 2017; Chen et al., 2019; Pechenik, 1999; Shikuma et al., 2014, 2016; Vlisidou et al., 2019). The close relatedness of BIS with other structures promoting microbe-eukaryotic interactions (MACs, Afp, PVCs and Subtype-4 T6SS) suggests that BIS mediate interactions between bacterial species within the human microbiome or Bacteroidales bacteria and their human host.

If BIS do interact with human cells, they may promote either symbiotic or pathogenic interactions. Injection systems closely related to BIS are described to mediate both beneficial and infectious microbe-host relationships. For example, MACs mediate metamorphosis of a marine tube worms (Ericson et al., 2019; Shikuma et al., 2014) and a Subtype-4 T6SS (Böck et al., 2017) mediates membrane interaction between *A. asiaticus* and its amoeboid host. In contrast, Afp and PVCs inject toxic effectors into insects (Hurst et al., 2004; Yang et al., 2006). While a Subtype-3 T6SS has been described in *Bacteroides* bacteria, we speculate that BIS could be the first Bacteroidales CIS to mediate microbe-animal interactions.

Structures that are closely related to BIS have been described to form T6SS that act from within a bacterial cell (e.g. Subtype-4 T6SS from *A. asiaticus*; Böck et al., 2017) or eCIS that are released by cell lysis (e.g. MACs and PVCs; (Shikuma et al., 2014; Vlisidou et al., 2019). While we currently do not know whether BIS generate a T6SS and/or an eCIS structure, we do know that intestinal bacteria like *Bacteroides* are physically separated from the intestinal epithelium by a layer of mucus (Marcobal et al., 2011; Tropini et al., 2017; Van Der Waaij et al., 2005). In contrast to T6SS that are bound to and act from within bacterial cells, an eCIS from Bacteroidales could cross this mucus barrier to inject effectors into human epithelial cells.

We show here that BIS are produced by Bacteroidales *in vivo* during colonization of humanized mice (Ridaura et al., 2013) and under laboratory conditions with various carbon sources (McNulty et al., 2013). However, we do not yet know the conditions that promote BIS production within normal or diseased human individuals. BIS may not have been extensively described before this work because they likely evolved independently from previously described CIS like Subtype-3 T6SS (Böck et al., 2017; Coyne et al., 2016) and possess significantly divergent sequence homologies (**Figure 1B**). Like other described CIS, BIS gene clusters possess genes encoding the syringe-like structural components and could encode for effectors that elicit specific cellular responses from target host cells. For example, the closely related injection system called MACs possesses two different effectors; one effector protein promotes the metamorphic development of a tubeworm (Ericson et al., 2019) and a second toxic effector kills insect and mammalian cell lines (Rocchi et al., 2019).

Many correlations between Bacteroidales abundances in the human gut and host health are currently unexplained. Future research into the conditions that promote the production BIS and its protein effectors will yield new insight into how Bacteroidales abundances correlate with host health, potentially by promoting the direct interaction between the microbiome and human host. In addition to their effect on host health, BIS provide the tantalizing potential as biotechnology platforms because they may be manipulated to inject engineered proteins of interest into other microbiome bacteria or directly into human cells.

## MATERIALS AND METHODS

### Phylogenetic analyses of CIS structural proteins

Whole genomes and assembled contigs of representative phage-like clusters (**Table S2**) were downloaded from NCBI to construct a database using BLAST+(2.6.0). The *B. cellulosilyticus* WH2 Sheath1 (WP 029427210.1) and Tube1 (WP 118435218) protein sequences were downloaded from NCBI and a tBLASTn was performed against the genome database. The recovered nucleotide sequences were then translated using EMBOSS Transeq (EMBL-EBI, https://www.ebi.ac.uk/Tools/st/emboss_transeq/) to generate a list of protein sequences. To capture highly divergent protein homologs, we used T6SS-Hcp, VipA/B; Phage- gp18, gp19 as reference proteins. The amino acid sequences were aligned using the online version of MAFFT (v7) with the iterative refinement alignment method E-INS-i. The aligned fasta file was converted into a phylip file using Seaview (Gouy et al., 2010). PhyML was performed through the ATCG Bioinformatics web server and utilized the Smart Model Selection (SMS) feature and the Maximum-Likelihood method (Guindon et al., 2010; Lefort et al., 2017). The best models for the Sheath1 and Tube1 phylogenies were rtREV+G+F and LG+G+I+F, respectively. Bootstrap values (1000 resamples) were calculated to ensure tree robustness. The Maximum likelihood tree topology was confirmed using other methods, including Neighbor joining, Maximum-Parsimony, Minimum-Evolution, and UPGMA. Trees were manipulated and viewed in iTOL (Letunic and Bork, 2016).

### BIS gene cluster synteny analyses

To identify CIS gene clusters in Bacteroidetes we used a modified protocol used to identify T6SS (Coyne et al., 2016). Briefly, the assemblies for 759 *Bacteroides* and *Parabacteroides* genomes included in the Refseq database (release 92, 26553804) were downloaded. Proteins from assembly were searched with HHMER v3.2.1 (http://hmmer.org/) for a match above the gathering threshold of Pfam HMM profile ‘phage_sheath_1’ (PF04984) (El-Gebali et al., 2019). For each match, up to 20 proteins were extracted from either side. All proteins from the resulting set (phage sheath + 20 proteins) were sorted by length and clustered at 50% amino acid identity using UClust v1.2.22q (Edgar, 2010). Clusters containing ≥ 4 members were analyzed further. Cluster representatives were annotated using protein-profile searches against three databases: the Pfam-A database using HMMER3 (El-Gebali et al., 2019), the NCBI Conserved Domain Database using RPS-BLAST (Marchler-Bauer et al., 2017, 2011; Marchler-Bauer and Bryant, 2004), and the Uniprot30 database (accessed February 2019, available from http://www.user.gwdg.de/~compbiol/data/hhsuite/databases/hhsuite_dbs/) using HHblits (McGarvey et al., 2019; Remmert et al., 2012). Multiple sequence alignments were automatically generated from three iterations of the HHblits search and used for profile-profile comparisons against the PDB70 database (accessed February 2019, available from http://www.user.gwdg.de/~compbiol/data/hhsuite/databases/hhsuite_dbs/). Significant hits to cluster representatives were used to assign an annotation to all proteins contained within the parent cluster. Manual inspection of *Bacteroides* and *Parabacteroides* loci enabled consistent trimming of each genetic architecture; specifically, the genes intervening DUF4255 and FtsH/ATPase were retained.

### Metagenomic mining analyses

To find the prevalence of the BIS genes within the Human Microbiome Project, we downloaded 11219 metagenomes using NCBI’s fastq-dump API. The metagenomes were parsed where: left-right tags were clipped, technical reads (adapters, primers, barcodes, etc) were dropped, low quality reads were dropped, and paired reads were treated as two distinct reads. A subject database was created from the amino-acid sequences of the 18 BIS genes. Then the fastq files were piped through seqtk (Shen et al., 2016), to convert them to fasta format, which was then piped to DIAMOND via stdin. Then DIAMOND aligned the six-frame translation of the input reads against the subject database, with all default parameters. For each metagenome the number of non-mutually exclusive hits to each CIS gene were then summed providing a hit ‘count score’. From the hit counts, a heatmap was created by taking the number of hits of each gene per metagenome and dividing by the total number of reads and multiplying the result by one million, which was then log10(x+1) transformed. To estimate the co-occurrence between pairs of genes, the previous the hit count scores from the previous calculation were taken, and for each pair combination, the hit count of the lower of the two genes was added to a running total. The co-occurrence was then visualized on a network graph, where each edge corresponds to the number of times the pair of genes co-occurred in all the metagenomes (Csardi and Nepusz, 2006; Hadley, 2016) (R core Team 2017 - https://www.R-project.org/; ggraph - https://CRAN.R-project.org/package=ggraph)

### Metatranscriptome mining analyses

Fastq files from transcriptomes were downloaded from the Sequence Read Archive using the sra toolkit (https://www.ncbi.nlm.nih.gov/sra/docs/toolkitsoft/). Low quality reads were removed using prinseq++ (https://peerj.com/preprints/27553/). Reads were compared to the amino acid sequences of the *Bacteroides cellulosilyticus* WH2 BIS protein cluster using blastX with an E-value cutoff of 0.001. The best hit for each read was kept. Hits to each protein were normalized by the number of reads of each transcriptome and the length of each protein using the program Fragment Recruitment Assembly Purification (https://github.com/yinacobian/frap).

## ACKNOWLEDGEMENTS

We thank Dr. Martin Pilhofer for providing constructive comments on the manuscript. This work was supported by the Office of Naval Research (N00014-17-1-2677, N.J.S. and S.B.), Office of Naval Research (N00014-16-1-2135, N.J.S), Alfred P. Sloan Foundation, Sloan Research Fellowship (N.J.S.), San Diego State University (N.J.S.), National Science Foundation PIRE grant (OISE1243541, F.L.R.).

## SUPPLEMENTARY MATERIAL

**TABLE S1.**

|  | Protein | Locus tag | *B. cell.*<br>WH2 CIS<br>protein | *B. cell.* WH2<br>Locus tag | Coverage | e-value | Identity |
| --- | --- | --- | --- | --- | --- | --- | --- |
| *P. luteo* MACs | MacB | WP_029427202.1 | baseplate | WP_029427202.1 | 84% | 5.00E-32 | 22% |
| *C. hertigii* T6SS | baseplate | WP_014934193.1 |  |  | 85% | 2.00E-57 | 23% |
| *S. entomophila* Afp | baseplate | WP_010895813.1 |  |  | 69% | 7.00E-18 | 25% |
| *P. asymbiotica* PVCs | baseplate | WP_015834374.1 |  |  | 68% | 3.00E-16 | 25% |
| *A. asiaticus* T6SS<sup>iv</sup> | baseplate | WP_012472726.1 |  |  | 77% | 2.00E-67 | 24% |
| *P. luteo* MACs | MacS | WP_039609824.1 | Sheath1 | WP_029427210.1 | 99% | 6.00E-94 | 35% |
| *C. hertigii* T6SS | Sheath | WP_014934609.1 |  |  | 89% | 8.00E-65 | 50% |
| *S. entomophila* Afp | Sheath1 | WP_010895805.1 |  |  | 87% | 9.00E-49 | 45% |
| *P. asymbiotica* PVCs | Sheath1 | WP_015835470.1 |  |  | 77% | 4.00E-54 | 46% |
| *A. asiaticus* T6SS<sup>iv</sup> | Sheath | WP_012473177.1 |  |  | 91% | 5.00E-66 | 50% |
| *P. luteo* MACs | MacS | WP_039609824.1 | Sheath2 | WP_029427209.1 | 85% | 1.00E-63 | 52% |
| *C. hertigii* T6SS | Sheath | WP_014934609.1 |  |  | 97% | 7.00E-122 | 39% |
| *S. entomophila* Afp | Sheath2 | WP_010895804.1 |  |  | 94% | 1.00E-51 | 48% |
| *P. asymbiotica* PVCs | Sheath2 | WP_015835471.1 |  |  | 97% | 3.00E-54 | 47% |
| *A. asiaticus* T6SS<sup>iv</sup> | Sheath | WP_012473177.1 |  |  | 97% | 3.00E-114 | 40% |
| *P. luteo* MACs | MacT1 | WP_039609825.1 | Tube1 | WP_007212392.1 | 97% | 5.00E-23 | 35% |
| *C. hertigii* T6SS | Tube | WP_014934612.1 |  |  | 73% | 6.00E-05 | 21% |
| *S. entomophila* Afp | Tube | WP_010895803.1 |  |  | 90% | 7.00E-32 | 34% |
| *P. asymbiotica* PVCs | Tube1 | WP_015835472.1 |  |  | 95% | 4.00E-31 | 33% |
| *A. asiaticus* T6SS<sup>iv</sup> | Tube | WP_012473180.1 |  |  | 84% | 3.00E-08 | 22% |
| *P. luteo* MACs | MacT2 | WP_039609826.1 | Tube2 | WP_007212393.1 | 92% | 6.00E-01 | 25% |
| *C. hertigii* T6SS | Tube | WP_014934612.1 |  |  | 57% | 1.00E-03 | 24% |
| *S. entomophila* Afp | Tube | WP_010895803.1 |  |  | 77% | 9.00E-09 | 24% |
| *P. asymbiotica* PVCs | Tube2 | WP_015834383.1 |  |  | 93% | 2.00E-09 | 23% |
| *A. asiaticus* T6SS<sup>iv</sup> | Tube | WP_012473180.1 |  |  | 92% | 3.00E-09 | 26% |

**TABLE S2.**

| Species | Strain | Genome<br>Accession | Sheath Protein<br>Accession | Tube Protein<br>Accession |
| --- | --- | --- | --- | --- |
| *Candidatus Amoebophilus asiaticus* | 5a2 | CP001102.1 | WP_012473177.1 | WP_012473180.1 |
| *Bacteroides cellulosilyticus* | WH2 | CP012801.1 | WP_029427210.1 | WP_007212392.1 |
| *Bacteroides fragilis* | BFBE1.1<br>(BE1) | LN877293.1 | WP_005803145.1,<br>WP_053873779.1 | WP_005803146.1 |
| *Cardinium hertigii* | cEper1 | NC_018605.1 | WP_014934609.1 | WP_014934612.1 |
| *Enterobacteria phage P2* |  | NC_041848.1 | YP_009591452.1 | YP_009591453.1 |
| *Enterobacteria phage T4* |  | NC_000866.4 | NP_049780.1 | WP_015969329.1 |
| *Flavobacterium johnsoniae* | UW101 | NC_009441.1 | WP_012025137.1,<br>WP_012025251.1 | WP_012025138.1 |
| *Francisella tularensis subsp.*<br>*tularensis* | SCHU S4 | AJ749949.2 | WP_003023948.1 | WP_003022149.1 |
| *Parabacteroides sp.* | D25 | NZ_JH976500.1 | WP_009276514.1,<br>WP_008669225.1 | WP_005861441.1 |
| *Pseudoalteromonas luteoviolacea* | HI1 | KF724687.1 | WP_039609824.1 | WP_039609825.1 |
| *Pseudomonas aeruginosa* | PAO1 | NC_002516.2 | WP_003113197.1,<br>WP_003087596.1 | WP_003083317.1,<br>WP_003085175.1 |
| *Salmonella enterica subsp. enterica*<br>*serovar Typhi* | CT18 | NC_003198.1 | WP_000046142.1,<br>WP_000013884.1 | WP_000338756.1,<br>WP_001207653.1 |
| *Serratia entomophila (pADAP)* | A1MO2 | NC_002523.4 | WP_010895805.1 | WP_010895803.1 |
| *Vibrio cholerae O1 biovar El Tor* | N16961 | NC_002506.1 | WP_001882966.1 | WP_001142947.1 |
| *Photorhabdus asymbiotica* | ATCC43949 | NC_012962.1 | WP_015835470.1 | WP_015835472.1 |

**TABLE S3.**
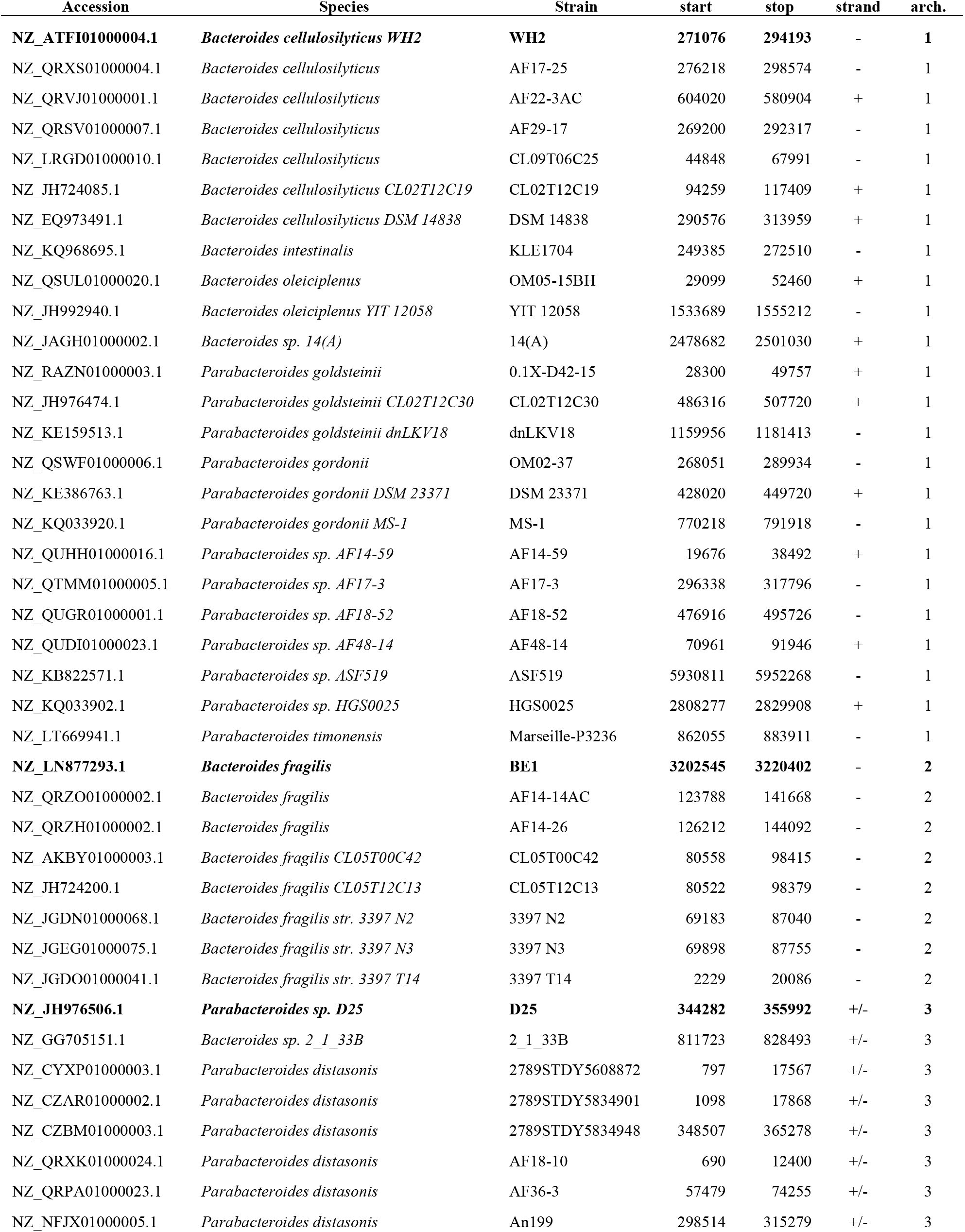
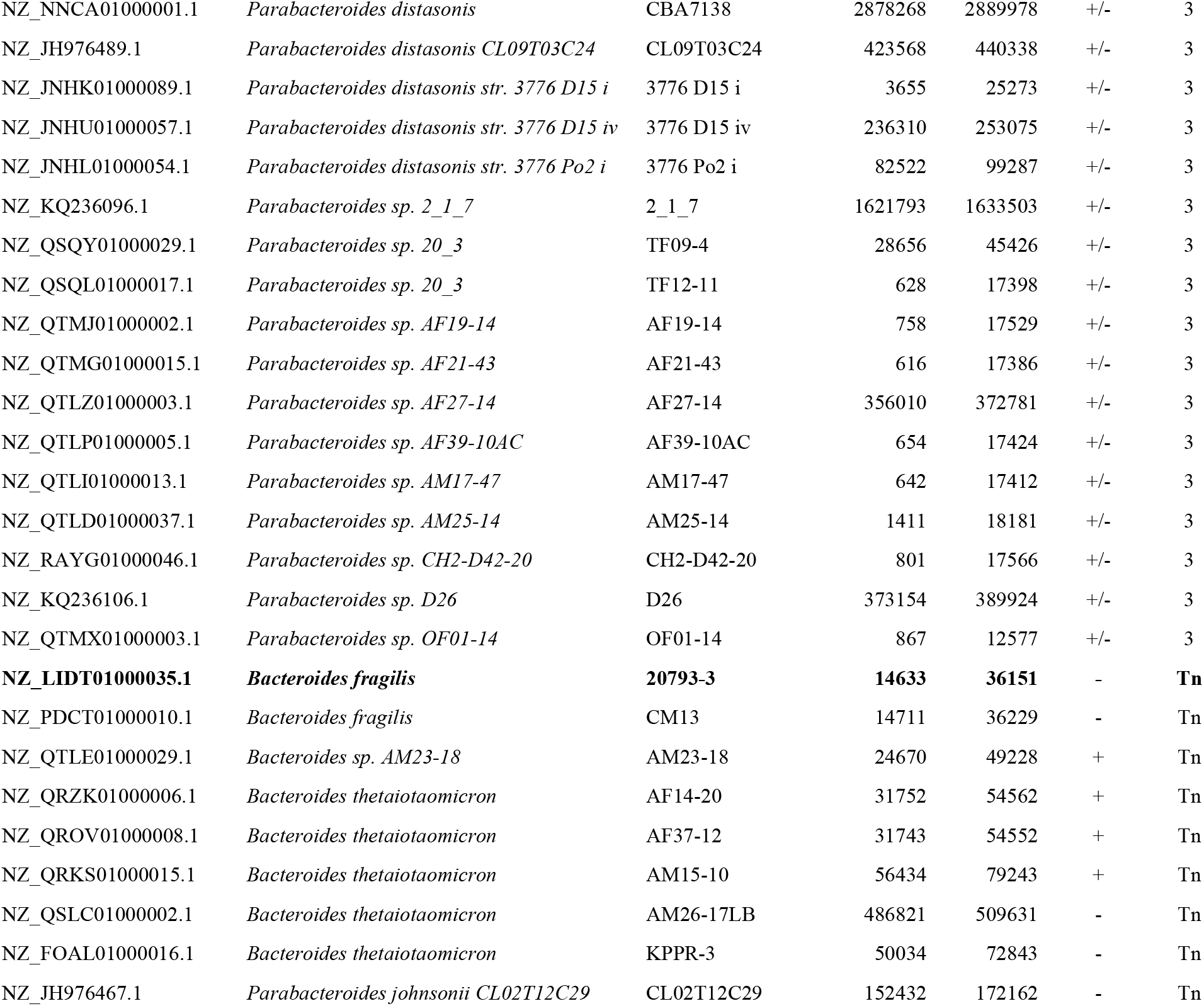

**Table S4.**

| Locus Tag (IMG:<br>GOLD Analysis<br>Project ID) | KEGG ID | Protein ID | Gene annotation | T4<br>homolog | pADAP<br>homolog |
| --- | --- | --- | --- | --- | --- |
| Ga0123724_112369 | BcellWH2_03967 | >WP_029427211.1 | DUF4255 | - | Afp16 |
| Ga0123724_112370 | BcellWH2_03968 | >WP_029427210.1 | Hypothetical protein | - | - |
| **Ga0123724_112371** | **BcellWH2_03969** | **>WP_029427209.1** | **Sheath1** | **gp18** | **Afp2/3/4** |
| **Ga0123724_112372** | **BcellWH2_03970** | **>WP_007212392.1** | **Sheath2** | **gp18** | **Afp2/3/4** |
| **Ga0123724_112373** | **BcellWH2_03971** | **>WP_007212393.1** | **Tube1** | **gp19** | **Afp1/5** |
| **Ga0123724_112374** | **BcellWH2_03972** | **>WP_007212394.1** | **Tube2** | **gp19** | **Afp1/5** |
| Ga0123724_112375 | BcellWH2_03973 | >WP_029427212.1 | Hypothetical protein | - | Afp6 |
| Ga0123724_112376 | BcellWH2_03974 | >WP_029427208.1 | LysM | gp6 | Afp7 |
| Ga0123724_112377 | BcellWH2_03975 | >WP_029427207.1 | Spike | gp5 | Afp8 |
| Ga0123724_112378 | BcellWH2_03976 | >WP_029427206.1 | Tip | gp5.4 | Afp10 |
| Ga0123724_112379 | BcellWH2_03977 | >WP_029427205.1 | Baseplate | gp25 | Afp9 |
| Ga0123724_112380 | BcellWH2_03978 | >WP_029427203.1 | Hypothetical protein | - | - |
| **Ga0123724_112381** | **BcellWH2_03979** | **>WP_029427202.1** | **Baseplate** | **gp27** | **Afp11** |
| Ga0123724_112382 | BcellWH2_03980 | >WP_029427201.1 | Hypothetical protein | - | Afp13 |
| Ga0123724_112383 | BcellWH2_03981 | >WP_029427200.1 | Baseplate | gp6 | Afp12 |
| Ga0123724_112384 | BcellWH2_03982 | >WP_029427199.1 | Hypothetical protein | - | Afp14 |
| Ga0123724_112385 | BcellWH2_03983 | >WP_029427198.1 | DUF4157 | - | - |
| Ga0123724_112386 | BcellWH2_03984 | >WP_007215181.1 | FtsH/ATPase | - | Afp15 |

**Figure S1.**
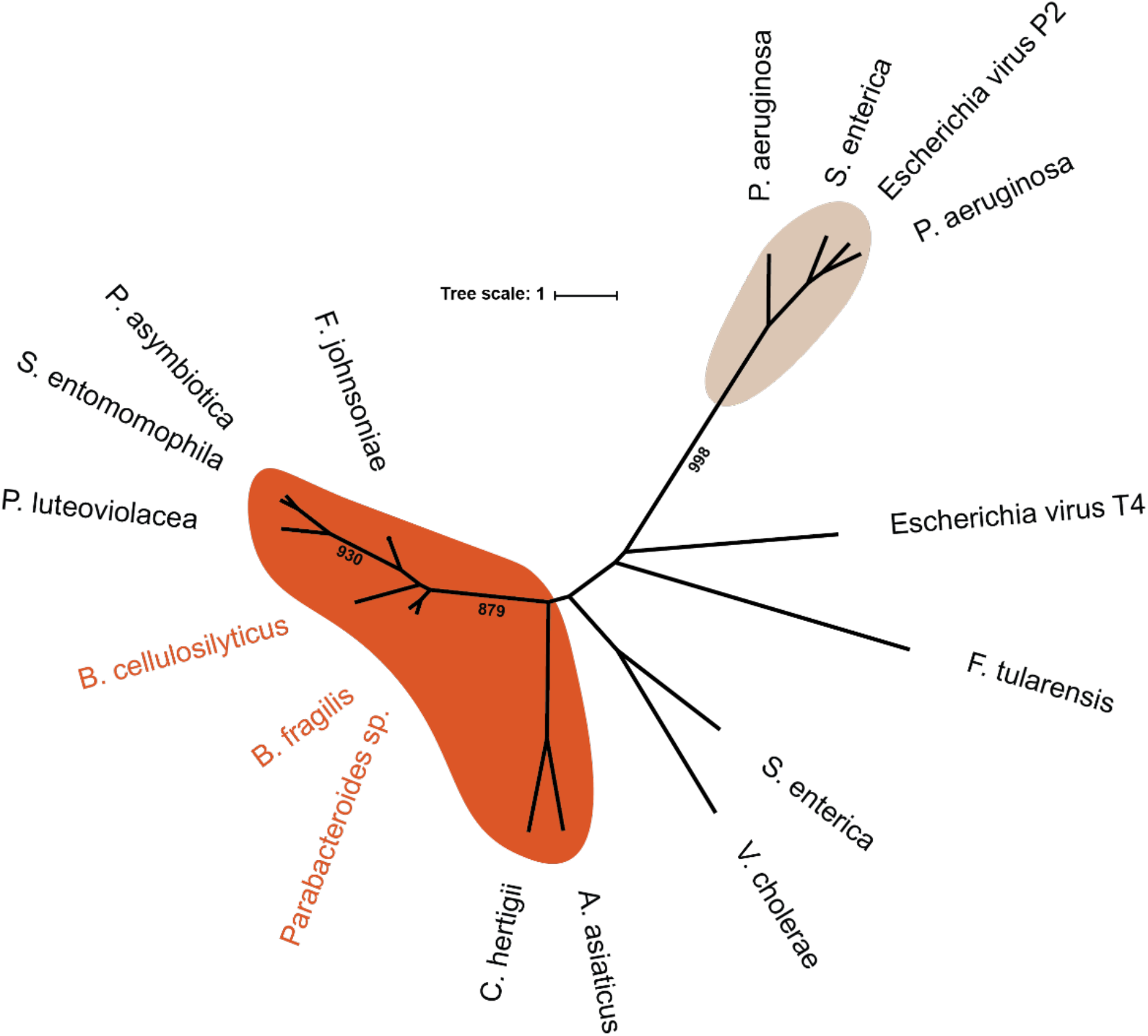
Unrooted phylogeny of CIS tube protein sequences. BIS group with known T6SS-iv and CIS (orange) and are distinct from known T6SS^i^ and T6SS^ii^ present in human pathogens, and known T6SS^iii^, characterized to mediate bacteria-bacteria interactions.

**Figure S2.**
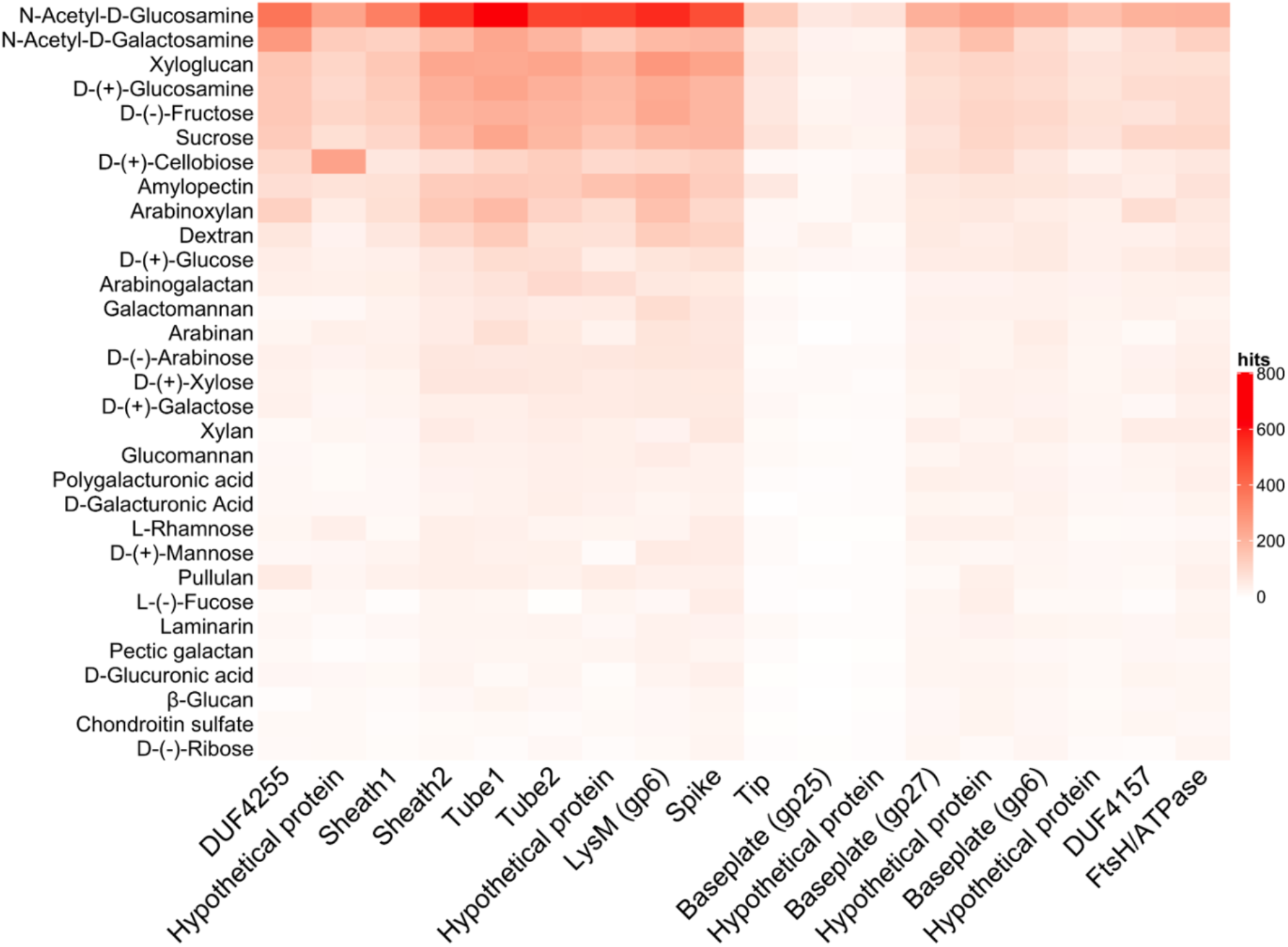
BIS genes are expressed during *in vitro* culture of *B. cellulosilyticus* WH2. Relative abundance of RNA hits to 18 major genes of the BIS in *B. cellulosilyticus* WH2 culture in MM supplemented with 31 different simple and complex sugars.

